# Induced-fit of the peptidyl-transferase center of the ribosome and conformational freedom of the esterified amino acids

**DOI:** 10.1101/032516

**Authors:** Jean Lehmann

**Affiliations:** Institute for Integrative Biology of the Cell (I2BC), CEA, CNRS, Université Paris-Sud, Campus Paris-Saclay, 91198 Gif-sur-Yvette, France

**Keywords:** Ribosome, peptidyl-transferase center, induced-fit, aminoacyl-tRNA

## Abstract

The catalytic site of most enzymes can essentially deal with only one substrate. In contrast, the ribosome is capable of polymerizing at a similar rate at least 20 different kinds of amino acids from aminoacyl-tRNA carriers while using just one catalytic site, the peptidyl-transferase center (PTC). An induced-fit mechanism has been uncovered in the PTC, but a possible connection between this mechanism and the uniform handling of the substrates has not been investigated. We present an analysis of published ribosome structures supporting the hypothesis that the inducedfit eliminates unreactive rotamers predominantly populated for some A-site aminoacyl esters before induction. We show that this hypothesis is fully consistent with the wealth of kinetic data obtained with these substrates. Our analysis reveals that induction constrains the amino acids into a reactive conformation in a side-chain independent manner. It allows us to highlight the rationale of the PTC structural organization, which confers to the ribosome the very unusual ability to handle large as well as small substrates.

## INTRODUCTION

An induced-fit (or conformational change) has been identified in the peptidyl-transferase center (PTC) of the ribosome, in which the binding of the 3’ acceptor arm of an A-site aminoacyl tRNA triggers a major rearrangement of two ribosome residues, U2506 and U2585 (*E. coli* numbering throughout this paper) (Schmeing et al. 2005a). The role of this induction in catalysis is however not clear. Upon induction, U2585 moves away from the reactive site in a way consistent with the hypothesis that it is protecting the peptidyl tRNA ester bond from premature hydrolysis in the uninduced state (Schmeing et al. 2005a). This interpretation was however questioned (Trobro and Åqvist 2006) on the grounds that puromycin (Pm) reacts at a very high rate (Monro and Marcker 1967; Sievers et al. 2004; Schroeder and Wolfenden 2007) despite the fact that this minimal substrate is unable to trigger the PTC conformational change. With Pm as the A-site substrate, the catalytic power of the ribosome has been attributed to an entropy reduction (as compared with a similar reaction in solution) (Sievers et al. 2004; Schroeder and Wolfenden 2007), a phenomenon that was proposed to result from the preorganization within the ribosome of a water molecule stabilizing the transition state (Trobro and Åqvist 2005). This explanation however also left the role of the induced-fit unclear. In order to establish the thermodynamics and the kinetics of peptide bond formation with full-length substrates, the accommodation step (during which the 3’ acceptor arm of an A-site tRNA moves into the PTC) must be excluded from the chemical step, a (still unresolved) difficulty at the origin of some controversy in the literature (Bieling et al. 2006; Johansson et al. 2008; Ledoux and Uhlenbeck 2008; Wohlgemuth et al. 2008; Rodnina 2013). Experiments with Phe-tRNA^Phe^ show that the thermodynamic parameters are similar to those obtained with Pm despite the inclusion of the accommodation step into the overall peptidyl transfer reaction (Johansson et al. 2008). This is consistent with the result that the induced-fit is not critical with Pm. The methyl tyrosine side-chain of this minimal substrate is very similar to that of phenylalanine, an aspect that is central to our analysis. Notwithstanding some uncertainties associated with the accommodation step, key experimental facts have been established: (**1**) In the early 70s, Rychlík and colleagues (Rychlík et al. 1969; Rychlík et al. 1970) showed that the activity (i.e. the ability to promote peptide transfer) of minimal A-site acceptor substrates in the form of 2’(3’)-O-aminoacyladenosine (A-aa) is extremely dependent on the side-chain of the amino acids. The least active acceptor is consistently A-Gly, while A-Phe is always the best acceptor, with an activity comparable to that of Pm. These authors also established that the relative activity of the acceptors is affected by the nature of the donor substrate. (**2**) More recent experiments showed that the nature of the C-terminal amino acid of peptidyl-tRNA donors modulates the rate of peptide bond formation when Pm is used as an acceptor substrate (Muto and Ito 2008; Wohlgemuth et al. 2008). (**3**) When full-length tRNA is used on both A- and P-sites, the rate of peptide bond formation seems to be independent of the nature of the amino acid at physiological pH (Ledoux and Uhlenbeck 2008; Wohlgemuth et al. 2008), with the exception of proline (Pavlov et al. 2009). In addition, short peptides are known to interfere with the PTC activity owing to their interaction with the exit tunnel (Ramu et al. 2011).

The purpose of this paper is to present a model that accounts for the above experimental facts and fully explains the role of the induced-fit of the PTC. Our investigation was motivated by an earlier analysis predicting a significant influence of the amino acids side-chain on the kinetics of peptide bond formation in a context without (elaborate) catalytic site (Lehmann 2000). Because most kinetic studies on the ribosome do not show any substantial side-chain effect in normal condition, we sought to relate the induced-fit mechanism of the PTC with the ribosome’s management of the side-chains, leading to the observed kinetics standardization. A key element of our analysis comes from the observation that while the kinetics of peptide bond formation with minimal A-site substrates is strongly side-chain dependent, their common feature (a single nucleotide moiety) *prevents them from triggering PTC induction*. We thus examined the possibility that the structure of the PTC cavity in the uninduced state may accommodate and stabilize unreactive rotamers of A-aa substrates. In Section 1, the activity of these minimal substrates is analyzed in terms of reaction rate (*k_cat_*) and binding (*Km*). We show that these activities reflect a *k_cat_* dispersion among some of them. Following an overview of structural features characterizing the uninduced and induced states (Section 2), we show that the room available inside the PTC cavity and its flexibility in the uninduced state leaves some conformational freedom to the esterified amino acids (Section 3). This feature enables the stabilization of rotamers for which the amino group is not oriented for nucleophilic attack, thus explaining why some minimal substrates react poorly. In Section 4, we show that induction corresponds to a compaction of the PTC, which forces any natural L-aminoacyl ester to adopt a unique (reactive) conformation. On the whole, our results allow us to highlight the rationale of the PTC structure-function relationship, this catalytic site having the very unusual requirement to accommodate large as well as small substrates. It appears that the uninduced state is required to let large amino acids enter the catalytic site, in which they are readily positioned for nucleophilic attack (except in very specific cases), while induction is crucial for canceling the rotational freedom of some smaller amino acids, the most critical one being glycine. These effects went unnoticed in most ribosome studies because they are usually performed with Pm and Phe-tRNA^Phe^ as model substrates, with which these conformational effects might not be detectable.

## RESULTS & ANALYSIS

### 1. Rates of peptide bond formation with minimal A-site substrates

The nature of the side-chain of A-aa acceptor substrates has been shown to strongly affect the acceptor activity in the peptidyl transfer reaction on the ribosome (Nathans and Neidle 1963; Rychlík et al. 1969; Rychlík et al. 1970) (Fig. 1). This activity is furthermore affected by the nature of the P-site donor. The most efficient (A-Phe) and the least active (A-Gly, A-(D)Phe) acceptors are however the same in all three donor configurations tested *in* (Rychlík et al. 1970). Because the acceptor activity comprises *Km* and *k_cat_*, catalytic rate constants cannot be extrapolated from the data of Figure 1, and this information could not be retrieved from more recent publications. There are however clear indications that the huge dispersion in acceptor activity predominantly reflects a variability of *k_cat_* among some substrates. Considering glycine and alanine (Fig. 1), the ratio of the slopes at the origin reveals a ~500-fold difference in acceptor activity. Yet the binding contribution of a methyl group does not exceed 1.3 kcal mol^−1^ (Hopkin 2008), while a more realistic value is closer to 0.9 kcal mol^−1^ (Plaxco and Goddard III 1994). The corresponding *K_d_* decrease is ~10-fold and 5-fold, respectively, well below the observed difference. Furthermore, the progression in acceptor activity (A-Phe, A-Ala, A-Leu, A-Ser, A-Met, …) does not follow a coherent affinity ranking (based on hydrophobicity). Thus, it can confidently be inferred that the various acceptor activities reflect differences in acceptor *k_cat_* values, at least among some of them. This issue is further examined in the discussion Section. A-Phe has approximately the same acceptor activity as Pm (Nathans and Neidle 1963; Rychlík et al. 1970), for which *k_cat_* has independently been established to ~5 s^−1^ with Met-Phe-tRNA^Phe^ as the donor substrate, under standard conditions (Sievers et al. 2004).

**Figure 1.**
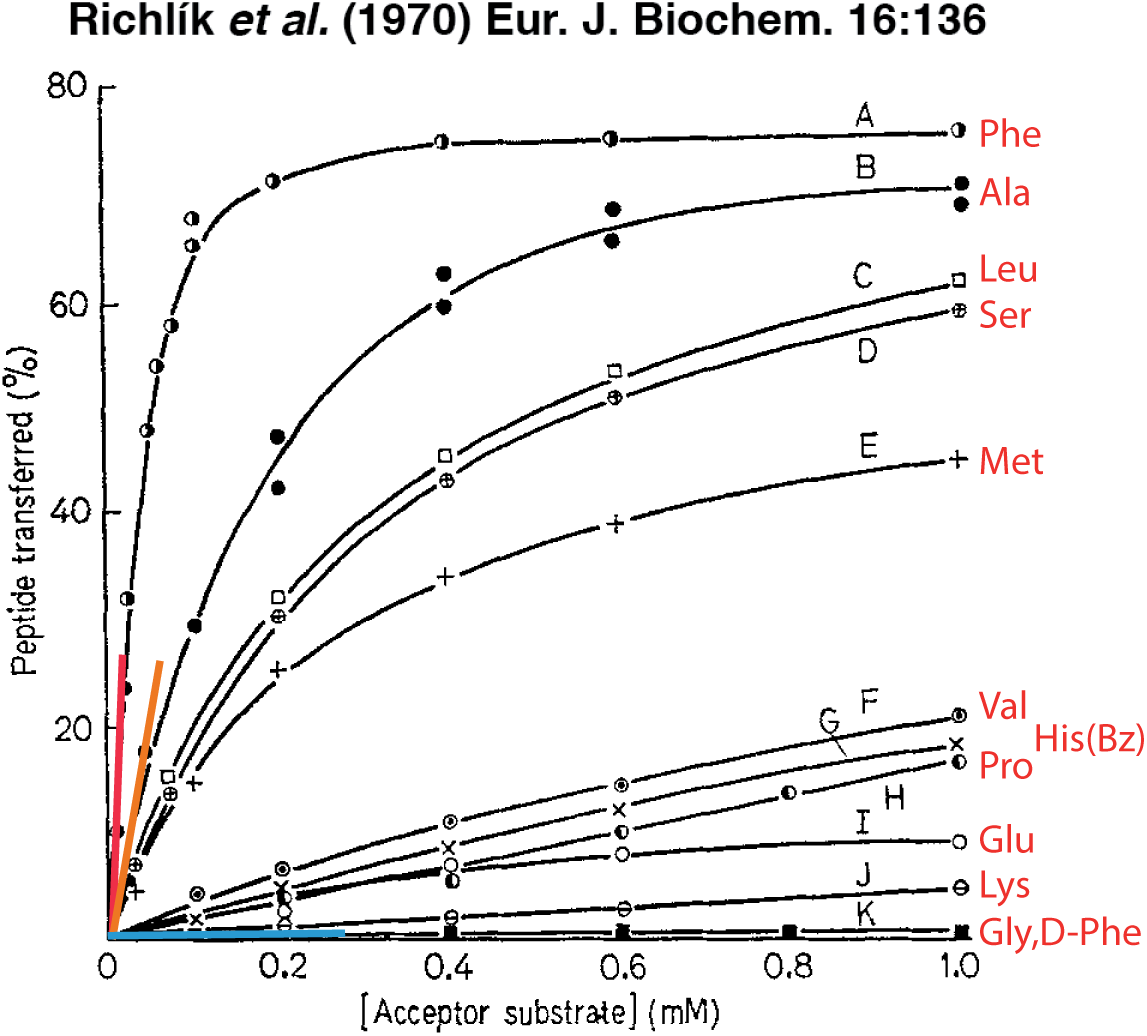
Transfer of peptide residues from (Lys)_n_-tRNA to minimal acceptor substrates in the form of 2’(3’)-O-aminoacyladenosine (A-aa). The relative values of the slopes shown at the origin are ~19 (A-Phe, in red), ~5.2 (A-Ala, in orange) and ~0.01 (A-Gly and A-(D)Phe, in blue). For each acceptor concentration, the amount of peptide transferred (in %) was determined after a 40-min incubation at 35°C (see the original publication for details). Adapted from (Richlík et al. 1970). Reproduced with kind permission of John Wiley & Sons, Inc.

In a more recent series of experiments, the influence of various C-terminal amino acids of P-site peptidyl donors on the kinetic constant was determined with Pm as the A-site acceptor (Wohlgemuth et al. 2008). In that case, the amplitude of the side-chain effect is less, of about one order of magnitude (*k_cat_* ~10–100 s^−1^), with the notable exception of proline (*k_cat_* ~0,1 s^−1^).

So far, no structural explanation has been proposed for these side-chain effects, observed when the (A-site) acceptor is a minimal substrate (Pm or A-aa). They were originally thought to result from various A-site binding propensities (Nathans and Neidle 1963; Rychlík et al. 1970) or from perturbed positioning owing to the small size of these substrates (Wohlgemuth et al. 2008). The most remarkable fact is that they are not observed when A-site and P-site substrates consist of full-length aminoacyl-tRNA and peptidyl-tRNA, respectively (Ledoux and Uhlenbeck 2008; Wohlgemuth et al. 2008). Although, in this configuration, the peptidyl-transfer reaction exhibits a measurable pH-dependence that is different for each amino acid (Johansson et al. 2011), the *pK_a_* contribution to *k_pep_* (a rate constant encompassing tRNA accommodation and peptidyl transfer) is comparatively small at physiological pH: *k_pep_* spans from ~6.0 to ~27 s^−1^ at pH 7.5 (Johansson et al. 2011).

The adenosine moiety of minimal substrates is known to form an A-minor interaction with G2583 (Nissen et al. 2001; Bashan et al. 2004; Voorhees et al. 2009). Because the side-chain of some of these substrates may clearly not modulate this interaction to an extent compatible with their different activities (see above), we examined whether some low activities could be related with the stabilization within the PTC cavity of unreactive rotamer(s) of the aminoacyl moiety. Our results (Section 3) are presented following an overview of the induced-fit mechanism (Section 2).

### 2. Induced-fit of the PTC and nature of the substrates

An induced-fit mechanism has been identified in the PTC of the ribosome by Steitz and coworkers (Schmeing et al. 2005a) (Figure 2). This mechanism could be outlined as follow: crystal structures show that in the absence of an A-site substrate, or when there is a minimal substrate no larger than CPm (see however discussion *in* Schmeing et al. 2005a, p. 523), the PTC remains in an uninduced state. In this state, U2506 forms a wobble base pair with G2583 and U2585 is pointing towards the A-site while being fairly mobile according to molecular dynamic simulations (Trobro and Åqvist 2006). When a minimal substrate is present, U2585(O4) however forms a hydrogen bond with its 2’-OH terminal ribose. When the A-site substrate is either CCPm or the 3’ end of a tRNA, the base triplet C_1_C_2_Pm or C_74_C_75_A_76_ squeezes in between U2555 (on which C_1/74_ stacks) and G2583 (with which Pm/A_76_ forms an A minor interaction) (Schmeing et al. 2005a; Voorhees et al. 2009). This compression is resolved through the breakage of the G2583-U2506 base pair, triggering U2506 to move to another equilibrium position that forces U2585 to stay away from the reaction center (Figure 2B).

**Figure 2.**
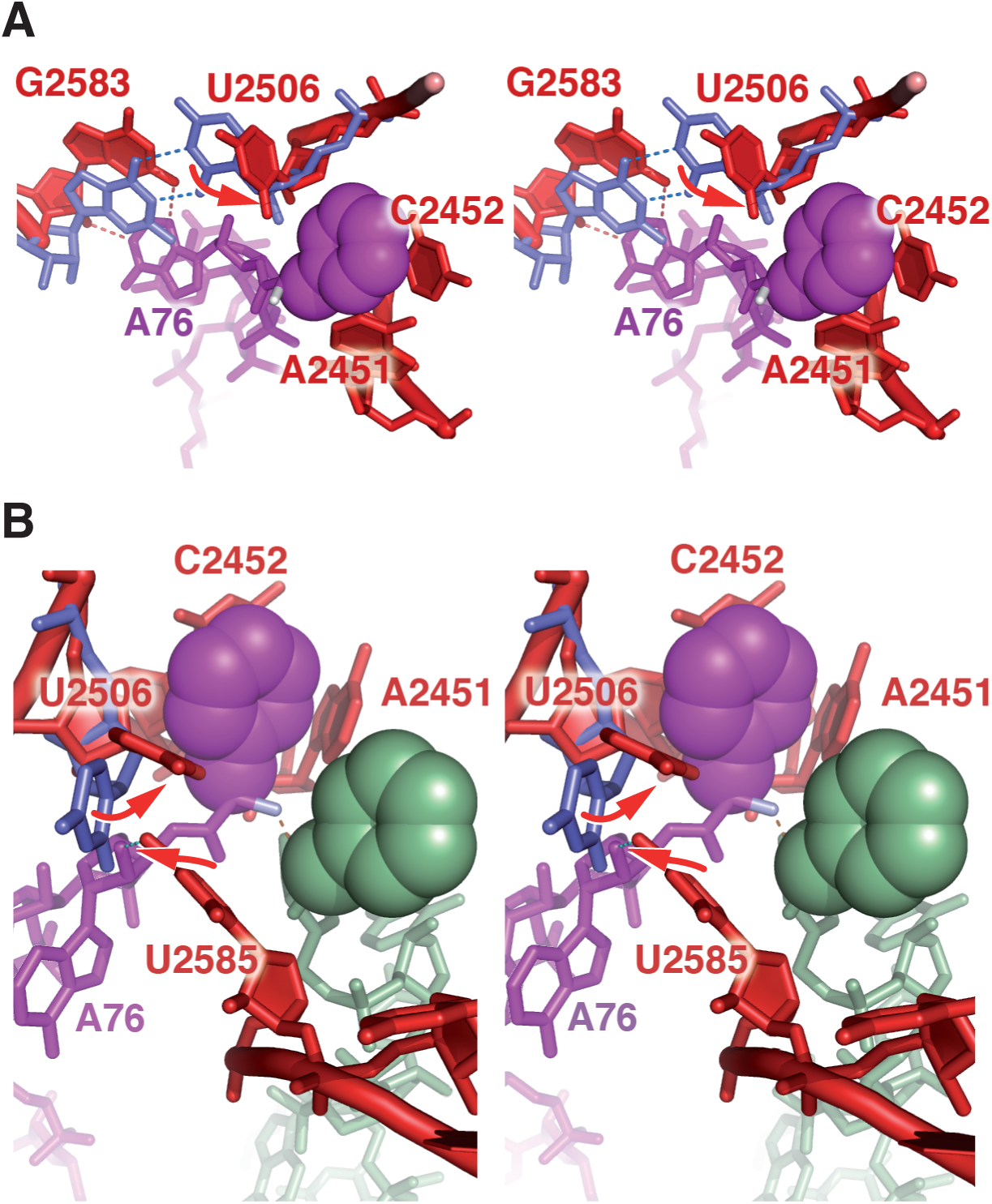
Induced-fit of the peptidyl-transferase center (Schmeing et al. 2005a): stereo views highlighting some major residues involved in the confinement of the A-site aminoacyl ester. (A) Following the binding of A76 to G2583 through an A-minor interaction, the G2583-U2506 wobble base pair breaks. As a result, U2506 moves (arrow) to a new equilibrium position, closer to the A-site aminoacyl ester. (B) Different view, showing the rearrangement of U2585 upon induction, which moves away from the reaction center. Ribosomal residues are shown in blue (before induction) and red (after induction). Residue U2585 is only shown after induction. A-site aminoacyl tRNA is in purple; P-site aminoacyl tRNA is in green. Phenylalanine side-chains are highlighted with van der Waals spheres. Residues in blue are from pdb 1VSA (Korostelev et al. 2006). Residues in red and aminoacyl tRNA are from pdb 2WDM and 2WDN (Voorhees et al. 2009). Superposition performed with *PyMOL*.

An examination of the two states made us realize that the resolution of the mentioned compression is responsible for a higher compactness of the PTC after induction, a so far unnoticed property that has a strong consequence for the conformational freedom of the esterifier amino acids. A comparison between uninduced and induced states indeed shows that there is more room for the A-site aminoacyl ester in the uninduced state, essentially because U2506 is kept away from the reaction center (Fig. 3).

**Figure 3.**
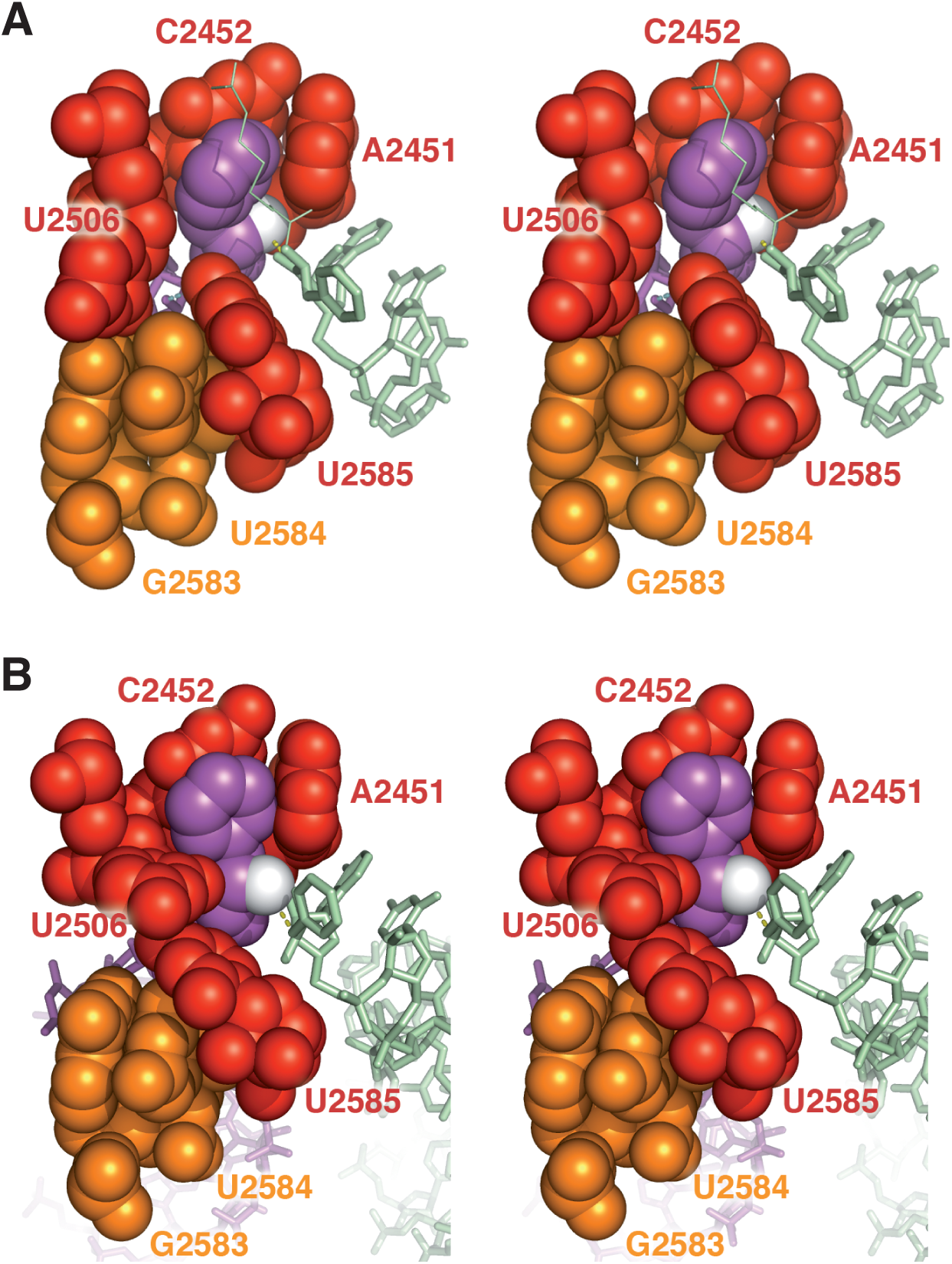
Peptidyl-transferase center before (A) and after (B) induction (stereo). (A) Uninduced state: when CA-Phe is on the A-site (in purple, aminoacyl ester highlighted with the van der Waals spheres) and CCA-Phe-biotin is on the P-site (in green, stick representation), no induced-fit is observed in the crystal (pdb 1VQ6, Schmeing et al. 200Sa). The biotin connector bound to the P-site aminoacyl ester in shown as thin lines. (B) Induced state: when full-length aminoacyl tRNA is on both A-site and P-site, an induced-fit is observed in the crystal (pdb 2WDM and 2WDN, Voorhees et al. 2009). In A and B, A-site Phe(N) (i.e. the nucleophile) is shown in white.

In the induced state, U2585 is squeezed in between U2584 and U2506, which keeps U2506 in close contact with the Cα atom of the amino acid (Fig. 3B). The side-chain lies within a cavity bounded by A2451, C2452 and U2506, with an opening towards the C-terminal peptidyl-tRNA. The aminoacyl ester is thus clearly locked in a conformation for which the amino group is properly positioned for nucleophilic attack. An analysis of the structural context of the reaction with the near-attack conformation (NAC) methodology (Lightstone and Bruice 1994, 1996, 1997, 1999; Griffin et al. 2012) reveals that with full-length substrates (pdb 2WDM and 2WDN, Voorhees et al. 2009), the amino group is at a distance *d* ~3.18 Å from the carbonyl carbon and at an angle of *α* ~35° to the normal of the carbonyl plane in the crystal. It is thus borderline of the defined NAC criteria, for which *d* is within 3.2 and 2.8 Å (when the nucleophile is the smaller oxygen) and *α ≤* 30° (Lightstone and Bruice 1994), although an uncertainty remains due to the resolution of the crystal structure (3.5 Å) and the possible influence of an O_ester_ → N_amide_ atom substitution. Assuming that confined molecules wedged in a conformation borderline NAC in the crystal fulfill the NAC criteria roughly half of the time (i.e. *p_NAC_* ~0.5), computational work predicts a relative rate constant of the order of 10^7^ M compared with free reactants in solution (Tables 1 and 2 *in* Lightstone and Bruice 1996).

In the uninduced state, the structural environment in the PTC cavity is less constraining for the aminoacyl ester (Fig. 3A). Because minimal substrates are unable to trigger the induced-fit, we inferred that the low activity observed with some of them had to be explained by the occurence of unreactive rotamer(s) of their aminoacyl moiety in that state.

### 3. The uninduced state of the PTC provides some conformational freedom to the aminoacyl esters

The minimal substrate A-Gly being the poorest acceptor (Fig. 1), it was used as a model to explore the possibility of unreactive rotamer(s) in the uninduced PTC cavity. With the 1VQ6 structure as a starting configuration, the A-site CPm side-chain was removed to get a molecule where the critical moiety is identical to A-Gly (Fig. 4A). Then, a plausible candidate obtained through a pi rotation of the free end around the *Ccarboxyl-Cα* bond (Psi dihedral angle) was immediately identified (Fig. 4B). This Psi(pi) rotamer has a total energy ~0.6 kcal mol^−1^ lower than the initial conformer (both *in vacuo* energy minimized), and readily fits the PTC cavity without any steric clash. In addition, the NH_2_ group of this rotamer is positioned for forming a hydrogen bond with U2585(O4) already in the (non energy minimized) 1VQ6 structure. In this configuration, U2585(O4) forms a bifurcated hydrogen bond with the 2’-OH terminal ribose and the NH_2_ group. An adjustment of the orientation of U2585 was performed to verify that both the distance and angle of the bifurcated interaction could be fully optimized. This was achieved while also replacing the P-site aminoacyl ester of the 1VQ6 structure, which is perturbed by a biotin, with the equivalent (underivatized) fragment of pdb 2WDM (see method Section). Computational works on bifurcated hydrogen bonds show that an energy of 3 to 4 kcal mol^−1^ is expected for the additional U2585(O4) - A-Gly(NH_2_) interaction (Sund et al. 2010; Feldblum et al. 2014). To establish this, it must be acknowledged that the NH_2_ group is > 4Å away from any potential H-acceptor in the initial configuration (Fig. 4A), the only possible candidate, A2451(N3), having been ruled out by mutational analysis (Erlacher et al. 2005). With a total energy difference of ~4.0 kcal mol^−1^, the A-Gly Psi(pi) rotamer is expected to be ~1000 times more populated than the reactive conformer. Because other (non NAC) Psi rotamers might also be significantly populated, this ratio constitutes a lower estimate to the reduction of the rate constant, as compared with amino acids trapped in the reactive conformation. We next investigated the possibility of the A-Phe Psi(pi) rotamer. A slight rotation of the phenylalaninyl moiety around the ester bond was required before energy minimization to avoid a clash with A2451 in the 1VQ6 structure. The rotamer resulting from this operation has a total energy ~1.2 kcal mol^−1^ lower than the initial molecule. In this situation, an orientation of U2585 corresponding to that of the (induced) 2WDN structure enables the phenylalanine side-chain to fit this region of the PTC cavity. This orientation of U2585 is plausible since it is observed in MD simulations even in the absence of strain (Trobro and Åqvist 2006). No steric clash is present in the resulting configuration (Fig. 5A), but the U2585(O4) - A-aa(NH_2_) hydrogen bond observed with A-Gly is prevented here due to the tilted orientation of U2585, a configuration that seems to be unstable. The original A-Phe conformer (Fig. 3A), for which the side-chain is buried inside the A2451/C2452 crevice, is thus expected to be more populated. Although our crude computational approach may not allow us to reliably assess the associated distribution, this model may still already account for the extreme cases of Figure 1, where the difference between the best acceptor (A-Phe) and the poorest acceptor (A-Gly) is of the order of 10^3^. The configuration of Figure 5A is expected to be favored in particular P-site contexts (see below).

**Figure 4.**
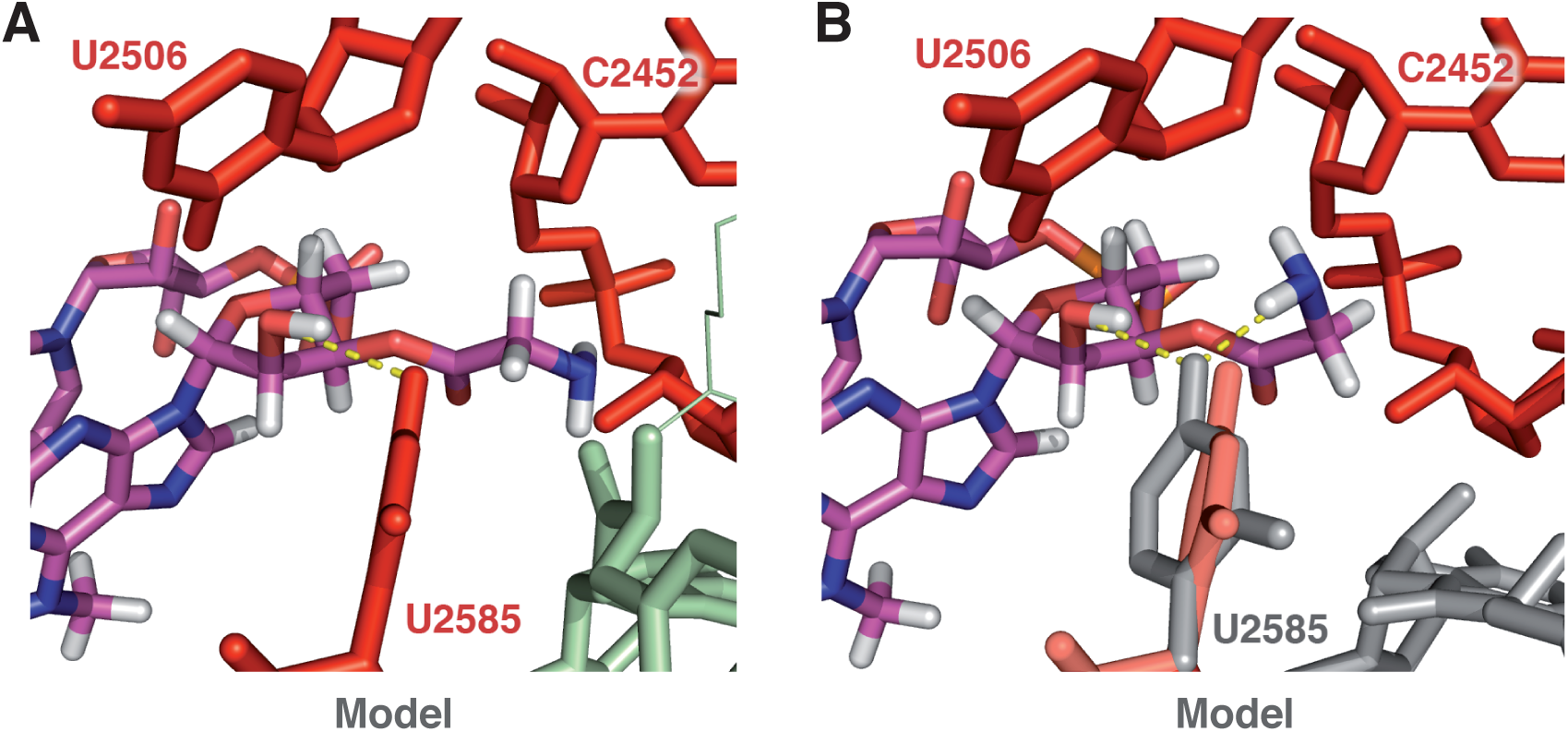
Energy minimized A-Gly 5’p fragments aligned in the peptidyl transferase center cavity (from pdb 1VQ6). (A) Initial conformer derived from the A-Phe 5’p fragment of the 1VQ6 structure. The ribose(2’-OH) – U2585(O4) hydrogen bond is shown in yellow. (B) Psi(pi) rotamer, enabling the additional Gly(NH_2_) - U2585(O4) hydrogen bond. The adjusted orientation of U2585 (in grey) results in both hydrogen bonds with a D-A distance of 3.0 Å. The P-site A-Phe fragment (in grey) is from pdb 2WDM (see method Section). Hydrogens (and atom colors) are only shown on the fragments.

**Figure 5.**
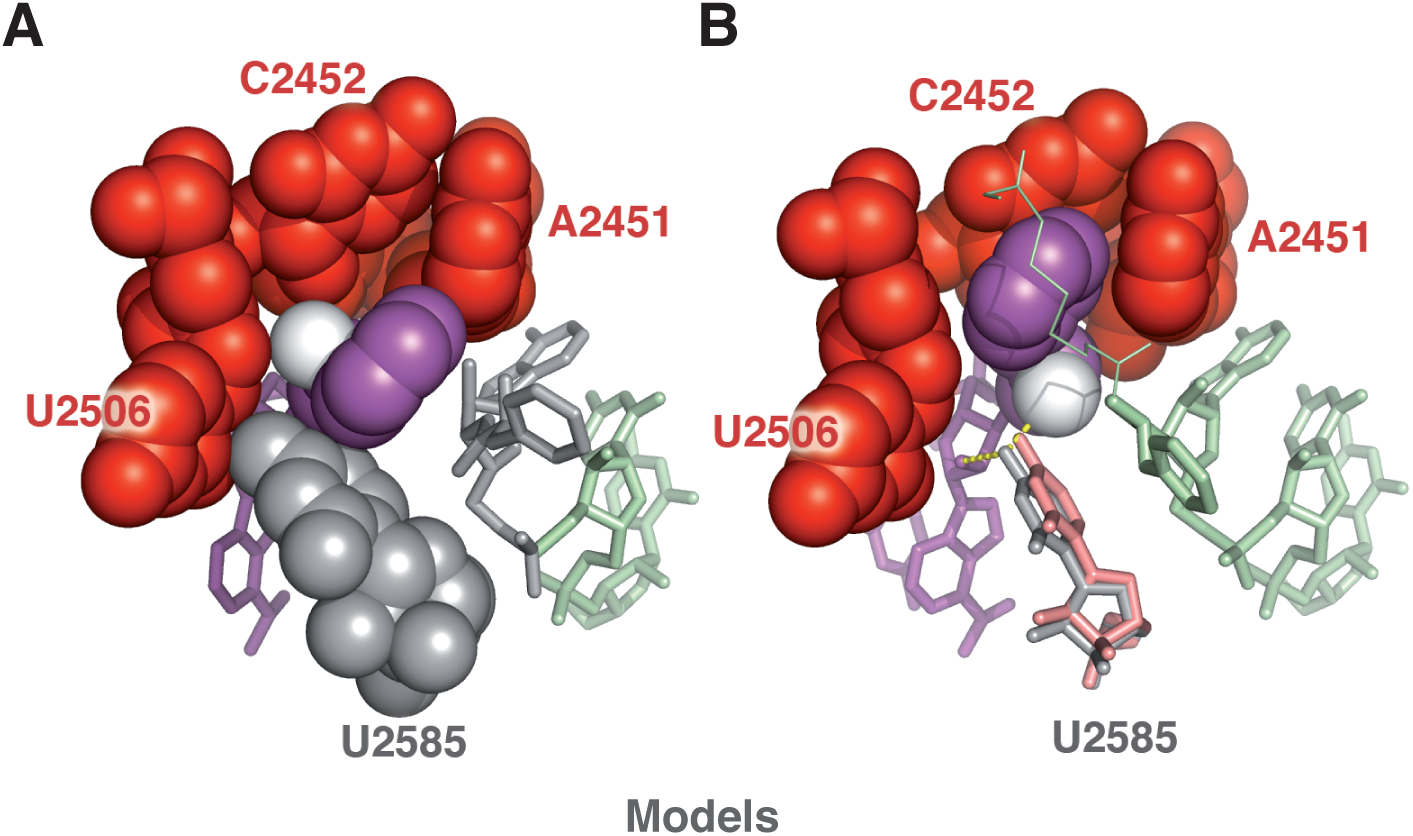
(A) Energy minimized Psi(pi) rotamer of A-Phe 5’p aligned in the PTC cavity (from pdb 1VQ6). This rotamer has a total energy ~1.2 kcal mol^−1^ lower than the original conformer. Residues in grey are from pdb 2WDM (P-site A-Phe) and pdb 2WDN (orientation of U2585). (B) A-(D)Phe 5’p in the PTC cavity (from pdb 1VQ6), built from the original phenylalanine ester of pdb 1VQ6 (see method Section). The position of U2585 (in grey) is slightly shifted from that of the 1VQ6 structure (in pale red) so as to potentialy allow a bifurcated hydrogen bond (in yellow). D-A lengths are both 2.9 Å; the DHA θ angle with Phe(NH_2_) is ~135°. In A and B, A-site Phe(N) is shown in white.

It is instructive to confront these structures with the situation provided by the D enantiomer of A-Phe, for which a total absence of reactivity was reported (Nathans and Neidle 1963; Rychlík et al. 1970). This molecule is indeed expected to keep its side-chain optimally positioned inside the A2451/C2452 crevice (Fig. 5B). Similarly to the situation observed with the A-Gly Psi(pi) rotamer (Fig. 4B), U2585(O4) may also form a bifurcated hydrogen bond, although the NH_2_ group is less optimally positioned due to a difference of ~(pi/3) in the Psi value. This unreactive configuration was already proposed to explain the prevalent exclusion of D-amino acids from translation (Zarivach et al. 2004; Agmon et al. 2004), which implies that induction usually goes wrong in that case (see below).

Another set of data accounted for by our model is the exceptionally low acceptor activity of Pm observed when the P-site C-terminal donor is proline (Muto and Ito 2008; Wohlgemuth et al. 2008). This configuration was examined while mutating the P-site phenylalanyl ester of Figure 5A to proline. It turns out that the Psi(pi) rotamer of A-Phe (or Pm) enables a π⋯CH interaction with proline on the P-site (supplementary Figure S1). This type of interaction is strong enough to occur within the aromatic(n)-proline(n+1) motif of proteins (Bhattacharyya and Chakrabarti 2003; Zondlo 2013); it is thus expected to stabilize the Pm Psi(pi) rotamer in that case. An examination of the structural context shows that geometric constraints prevent the occurrence of similar side-chain – side-chain interactions in the other amino acid configurations tested by Rodnina and coworkers (Wohlgemuth et al. 2008), e.g. Pm-Phe (see Fig. 5A). While a ~60 to 700-fold decrease in *k_cat_* is observed with proline on the P-site (Wohlgemuth et al. 2008), the mentioned π⋯CH interaction plausibly involves an energy of 1 to 3 kcal mol^−1^ (Morozov et al. 2004; Jovanovic et al. 2015), compatible with a ~100-fold change in the affinity ratio required to account for the observed *k_cat_* decrease.

### 4. The induced-fit orients the aminoacyl ester for nucleophilic attack

One justification of our hypothesis of unreactive Psi rotamers in the uninduced state is a stabilizing U2585(O4) - aa(NH_2_) hydrogen bond (Section 3). Induction is precisely removing the possibility of this interaction by moving U2585 away from the reaction center (Fig. 6). Furthermore, the NH_2_ group of any aa-tRNA Psi(pi) rotamer would clash with U2506(O2) (Fig. 6A), a destabilizing effect that disappears only with a full switch to the reactive conformation (Fig. 6B). This conformation is maintained because A76 is bound to G2583 while U2584 and U2585 are wedged in between G2583 and U2506 (Fig. 3B), and U2506 is locked in place by G2505 (supplementary Figure S2). This structural context suggests to us that the Cα atom of the aminoacyl ester is *held back* on the other side by A76(2’-OH) of the P-site tRNA, possibly in conjunction with A2451(2’-OH) (Lang et al. 2008). As a logical consequence, the 2’ hydroxyl group of A76 also contributes to orienting the NH_2_ group towards the carbonyl carbon (Fig. 6B), a possibility in line with the analysis of Green and coworkers (Zaher et al. 2011), and consistent with the significant drop in peptidyl transfer activity observed when this 2’-OH is replaced with 2’-H (Zaher et al. 2011).

**Figure 6.**
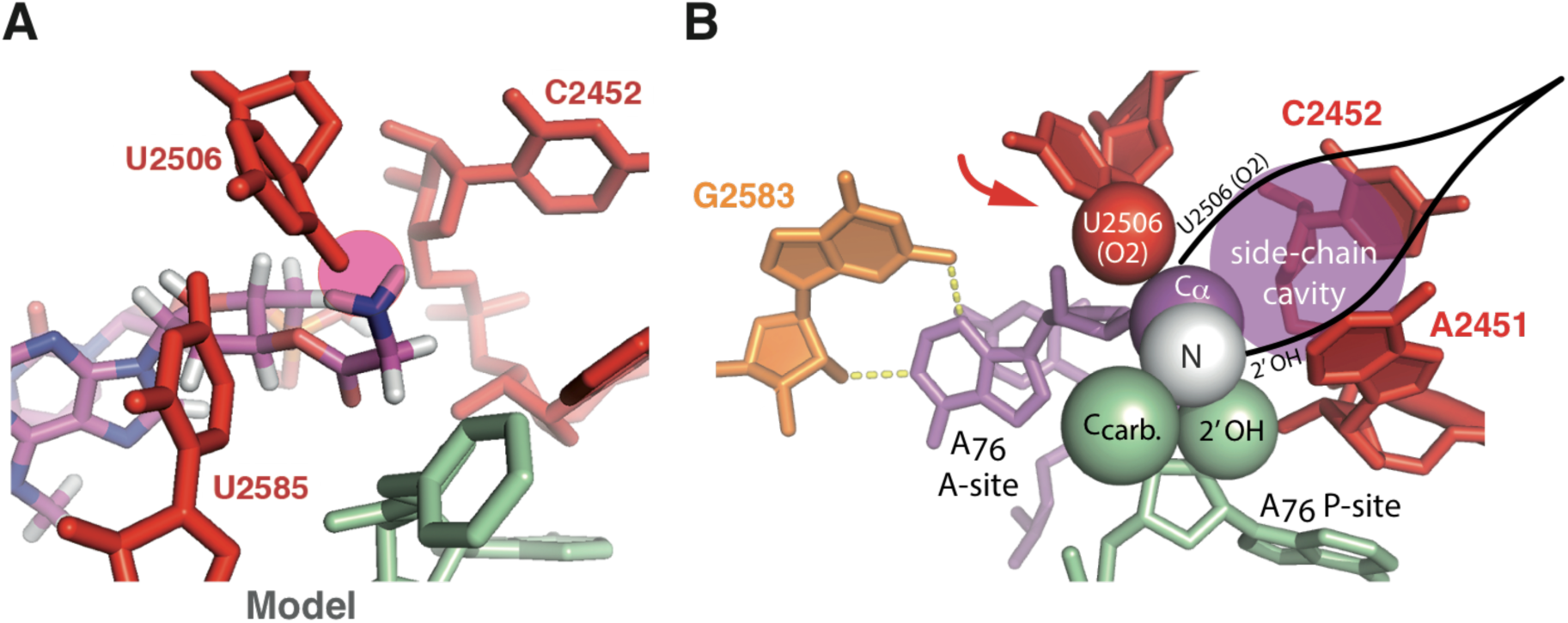
Structural constraints resulting from induction. (A) Energy minimized Psi(pi) rotamer of A-Gly 5’p hypothetically aligned in the peptidyl transferase center cavity in the induced state (from pdb 2WDN): a major clash of the amino group with U2506(O2) (pink circle) prevents the possibility of this rotamer (see also supplementary Figure S2). (B) Induced state: view highlighting the U2506(O2) – A76(2’-OH) Cα pinching mechanism enforcing the reactive conformation (the arrow symbolizes the swing of U2506 upon induction). The cavity by the C*α* atom accommodates L-amino acid side-chains. Key residues (or only key atoms, highlighted with van der Waals spheres) are shown. A-site aminoacyl tRNA is in purple, with Phe(N) in white (as in Fig. 3B); P-site aminoacyl tRNA is in green. Unmodified 2WDM and 2WDN structures (Voorhees et al. 2009).

The structural hallmark emerging from the above analysis is a mechanism in which U2506(O2) and A76(2’-OH) are pinching the C*α* atom upon induction, an action that orients the aminoacyl ester for nucleophilic attack. Remarkably, this mechanism preserves a cavity that is large enough to accommodate any (natural) L-amino–acid side-chain (Fig. 6B).

Despite the constraints resulting from induction, D-amino acids are incorporated only at very low yields (Fujino et al. 2013), a property resulting from the localization (with respect to the C*α* atom) of the mentioned cavity. Ribosome mutants selected for their ability to better incorporate D-amino acids are less efficient in wild-type protein synthesis (Table 2 *in* Dedkova et al. 2006). Furthermore, an analysis of ribosome mutants selected to incorporate β-amino acids suggests that PTC structures capable of better accommodating non α-L-amino acids are more flexible (Dedkova et al. 2012; Maini et al. 2015). Taken together, these observations suggest a positive correlation between compaction, D/L selectivity and catalytic efficiency.

## DISCUSSION

### Rationale of the peptidyl transferase center structure-function organization

Our analysis points out that the conformational freedom of some amino acid acceptors is at the origin of their low reactivity in the reaction of peptide bond when the ribosome is stuck in the uninduced state, which occurs when a minimal substrate is present on the A site. Why then does the PTC have an uninduced state? An already proposed explanation is that the orientation of U2585 in the induced state may not protect the P-site aminoacyl ester from premature hydrolysis as it does in the uninduced state (Schmeing et al. 2005a), a proposal supported by experiments with deacylated tRNA bound on the A-site, shown to promote such hydrolysis (Zavialov et al. 2002).^1^

A major role of the uninduced state identified by our study is to allow large amino acids enter the PTC cavity. Owing to its compactness, a pre-induced PTC would prevent these residues from going inside, a conclusion that can be drawn from the observation that U2506 is literally trapping their side-chain upon induction (supplementary Figure S3). Translation inhibition by Pm analogues with double-ring side-chains (L-Trp and *im*.-benzyl-L-His derivatives), for which particularly low efficiencies are observed (Harris et al. 1971), suggests that PTC penetration by very large amino acids may already be a problem in the uninduced state. This points out another role for the A-site tRNA’s 3’ acceptor arm binding, that helps push bulky residues inside the PTC cavity (a phenomenon already considered by Sharma et al. 2005). It also rationalizes the mobility of U2585 observed in MD simulations before induction (Trobro and Åqvist 2006), crucial to the entering of very large amino acids (supplementary Figure S3A). The low activity in peptidyl transfer reaction observed with the minimal form of these substrates (Section 1) thus most likely reflects a slow PTC penetration. The maximal activity observed with A-Phe and Pm indicates that these substrates optimally combine PTC penetration and reactivity inside the cavity (Vanin et al. 1974).

In brief, freezing the esterified amino acids into a reactive conformation by a compaction of the catalytic site requires an uninduced state to let *all the amino acids* enter the PTC.

### Why does the modern ribosome confine the esterified amino acids?

Measurements with full-length tRNA indicate that the kinetics of peptide bond formation is nearly uniform with natural aminoacyl ester substrates (Ledoux and Uhlenbeck 2008; Wohlgemuth et al. 2008). Our analysis reveals that this standardization is a consequence of a compaction of the PTC upon induction, that is remarkably not affected by the nature of the a-L-amino acid side-chains (Section 4). It can be mentioned that active site compaction by directed mutagenesis has been shown to increase the catalytic efficiency of an enzyme (Zhang and Klinman 2011).

A proper immobilization of the aminoacyl esters is essential for ensuring the highest possible rate. Studies on intramolecular reactions have long revealed that the *local* conformational freedom of a nucleophile has a huge impact on the associated rate constant (Beesley et al. 1915; Bruice and Pandit 1960; Milstien and Cohen 1970; Storm and Koshland 1970; Lightstone and Bruice 1997; Lightstone and Bruice 1999; Bruice and Benkovic 2000; Jung and Piizzi 2005; Kraut et al. 2003; Garcia-Viloca et al. 2004). Furthermore, both the position and size of carbon substituents close to the nucleophile are critical without confinement, bulky substituents typically conferring higher reaction rates to a considered intramolecular system. In the case^2^ of 3’ esterified amino acids, we are not aware of any data directly assessing a possible effect of the side-chains on the rate of peptide bond formation in the absence of confinement. However, in the context of a primitive kinetic scheme of translation, a correlation in the genetic code connecting the strength of the anticodon-codon association with the size of the amino acids was interpreted as a consequence of such side-chain effects (Lehmann 2000). Relating anticodon-codon stability to a dissociation rate constant *k_* and the size of the side-chains with *k_cat_* (large side-chains conferring the highest *k_cat_*), we interpreted the correlation as a situation for which the equality *k_cat_* ≈ *k_* is verified for all [amino acids │ codon] couples (Lehmann 2000; Lehmann et al. 2009), corresponding to an overall optimization of the rate of peptide bond formation on the early ribosome (additional considerations are discussed in supplementary Text S1). The possibility of this kinetic balancing was already suggested as early as 1985 (Remme and Villems 1985) (F. Michel, pers. comm.). The side-chain effect can explain the origin of the “trick” used by the modern ribosome to freeze any incoming α-L-amino acid ester into a same reactive conformation. Because the diversity of side-chains rules out the possibility of a direct interaction, the trick relies on a mechanism pinching the C*α* atom of the amino acids while preserving a large cavity capable of accommodating any natural side-chain (Fig. 3 & Fig. 6B).

### On the entropy of the activation energy

A seminal investigation (Sievers et al. 2004; Schroeder and Wolfenden 2007) has shown that the ribosome is essentially reducing the entropy of activation ΔS^‡^) of peptide bond formation as compared with an “equivalent” reaction in solution, a reduction that may account for the ~10^7^-fold rate enhancement produced by the ribosome (Sievers et al. 2004). An estimate we provide in Section 2 suggests that this effect may entirely result from substrate orientation, a possibility that was already supported (Schroeder and Wolfenden 2007; Moore and Steitz 2011). Substrate orientation is related to the thermodynamic concept of “ground state preorganization” (Armstrong and Amzel 2003; Storm and Koshland 1970; Lightstone and Bruice 1999; Bruice and Benkovic 2000; Kraut et al. 2003; Garcia-Viloca et al. 2004), in which the cost of binding a substrate with the right orientation for nucleophillic attack (an effect that is reducing ΔS^‡^) is paid through binding enthalpy.

An alternate (or complementary) possibility, claimed to explain the decrease of ΔS^‡^ compared with the reaction in solution, is an electrostatic preorganization of the active site (Trobro and Åqvist 2005). It was found in crystals and MD simulations that a water molecule trapped inside the ribosome may be involved in the stabilization of the oxyanion of the transition state (Trobro and Åqvist 2005; Schmeing et al. 2005b). Thermodynamics measurements (Sievers et al. 2004) show that ΔS^‡^ is not much lowered when the concentration of Pm is increased from a non-saturating to a saturating concentration, a result that was mentioned in support of the electrostatic preorganization effect (Trobro and Åqvist 2005). A saturating concentration however only results in the loss of (almost all) *translational* entropy, which does not imply that the substrate is already properly positioned in the active site to react (Storm and Koshland 1970). The alternate (or complementary) interpretation precisely states that the ΔS^‡^ reduction on the ribosome is due to ground state effects, in which substrate binding only occurs for conformers that are properly *oriented* to enter the reaction (Lightstone and Bruice 1999; Bruice and Benkovic 2000; Kraut et al. 2003; Garcia-Viloca et al. 2004), which implies the reduction of rotational and vibrational entropic terms. Although the present analysis may not directly assess the contribution of electrostatic preorganization (see Trobro and Åqvist 2005; Wallin and Åqvist 2010; Carrasco et al. 2011), it provides the evidence that ground state effects cannot be neglected.

## CONCLUSION

Our analysis reveals that the induced-fit of the PTC of the ribosome is required for the proper incorporation of *all* natural α-L-amino acids into a nascent protein. Depending on the A-site (aminoacyl) and P-site (C-terminal peptidyl) context, peptide bond formation may be prevented by the occurrence of Psi rotamer(s) of the A-site aminoacyl ester before induction. The concurrent binding of the tRNA’s 3’ acceptor arm triggers a compaction around the amino acid that forces it to adopt a reactive conformation. The induced-fit mechanism also prevents a premature hydrolysis during translation, a property identified in the seminal study of Schmeing et al. (2005a). Our results suggest that the rather large size of the ribosome structure around the PTC is due to architectural constraints required to establish a robust induced-fit mechanism. It may explain why the ribosome is a “versatile catalyst” (Rodnina 2013), capable of “packing” various substrates provide that they can properly undergo induction.

Our analysis centered on the structure and the origin of the genetic code (Lehmann 2000) predicted that the side-chains of the amino acids contributed to the establishment of the genetic code at the level of the ribosome in the absence of an elaborate PTC and a decoding center. In conjunction with optimizations at the level of tRNA (especially at the level of the anticodon loop), both sites have apparently evolved to free translation from basic physico-chemical constraints, resulting in a standardization and optimization of the kinetic constants. This standardization was likely required before higher levels of complexity in the genetic system (such as the regulation of gene expression) could further evolve.

## COMPUTATIONAL METHODS

A semi-quantitative assessment of the structures being investigated was performed using the *openbabel* package, version 2.3.2 (O’Boyle et al. 2011). 3’-O-aminoacyladenosine 5’ phosphate (A-aa 5’p) rotamers were *in vacuo* energy minimized with the steepest descent algorithm while using the MMFF94 force field. Convergence always occurred within 10,000 steps. Results are expressed in terms of an energy difference between rotamers. A-aa 5’p Psi(pi) rotamers were generated with the *obrotate* tool from the initial molecule of (or derived from) the 1VQ6 pdb file (Schmeing et al. 2005a), and subsequently energy minimized. Refined A-aa 5’p structures were aligned in the PTC cavity using the *PyMOL* alignment tool. Alignment was solely based on the adenosine moiety of the pdb file. U2585 is known to be a “universally mobile” residue of the PTC (Zarivach et al. 2004; Agmon et al. 2004; Trobro and Åqvist 2006). Depending on the context, the orientation of this nucleotide was adjusted following separately described procedures. Resulting configurations were checked for steric clash; additional energy contributions in the PTC were evaluated separately.

For the configuration with A-site A-Gly 5’p Psi(pi) rotamer (Fig. 4B), the P-site A-Phe fragment of pdb 2WDM (Voorhees et al. 2009) was aligned to that of pdb 1VQ6 (Schmeing et al. 2005a) solely based on the P-site A76 moiety. The orientation of U2585 was adjusted so as to obtain a bifurcated hydrogen bond in which each bond is characterized by a donor-acceptor (D-A) distance of 3.0 Å.

With the (A-site) A-Phe 5’p Psi(pi) rotamer (Fig. 5A), the P-site A-Phe fragment of pdb 2WDM has the same alignment as in Figure 4B. The orientation of U2585 was obtained from a *PyMOL* alignment of the 2580–2590 11-base fragment of 2WDN (Voorhees et al. 2009) (an induced structure) to that of 1VQ6. The A-site D enantiomer of A-Phe was built with *PyMOL* from the original A-site L-Phe residue of pdb 1VQ6 with a NH_2_ ↔ H permutation on the C*α* atom (Fig. 5B). The orientation of U2585 was slightly adjusted so as to position U2585(O4) in hydrogen bonding distance from Phe(NH_2_) while keeping hydrogen bonding potentiality with A76–2’OH, with no further optimization.

The Phe-Pro configuration (Fig. S1) was derived from Figure 5A. The P-site A-Phe fragment of 2WDM was mutated to A-Pro with the *PyMOL* mutagenesis tool, and a slight rotation (< 5°) of this residue around the ester bond was performed to prevent a steric clash with the A-site side-chain, with no further optimization.

A-site tryptophanyl ester molecules (Fig. S3) were obtained with the *PyMOL* mutagenesis tool from pdb 1VQ6 (uninduced structure) and pdb 2WDM and 2WDN (induced structure). No adjustment of PTC residues was performed.

The near attack conformation (NAC) methodology (Lightstone and Bruice 1994, 1996, 1997) was used to determine whether the conformer of the A-site phenylalanine ester of the (induced) 2WDM and 2WDN structure corresponds to (or is close to) a NAC conformer, i.e. a conformer that can perform nucleophilic attack. It requires the determination of the position of the nucleophile of the A-site phenylalanine ester with respect to the carbonyl carbon of the P-site phenylalanine ester, both in terms of distance to the carbonyl carbon (*d*) and angle to the normal of the carbonyl plane (α). This method has already been used to characterize conformations of reactive species in crystals (Griffin et al. 2012). The determination of *d* and *α* was achieved with *PyMOL*. The equation of the carbonyl plane in the crystal was determined from the *Ccarbonyl* and the connected O and C*α* atoms.

## ACKNOWLEDGEMENTS

I am very grateful to Daniel Gautheret for supporting my work in his laboratory, and for stimulating discussions about many aspects of RNA. I would like to thank François Michel and Benoît Masquida, who examined my analysis in detail, and provided many suggestions that improved the manuscript. Several aspects of this work were also improved thanks to comments and suggestions from Shixin Ye-Lehmann, Pradeep Kumar, Daniel Gautheret, Fabrice Leclerc, Thomas Simonson, Alexandre V. Morozov and Pamela Rodriguez. I also would like to thank Henri Grosjean for stimulating discussions about tRNA evolution and Pradeep Kumar for inspiring conversations in biological physics.

## SUPPORTING INFORMATION

Figures S1, S2 and S3; Text S1

**Figure S1.**
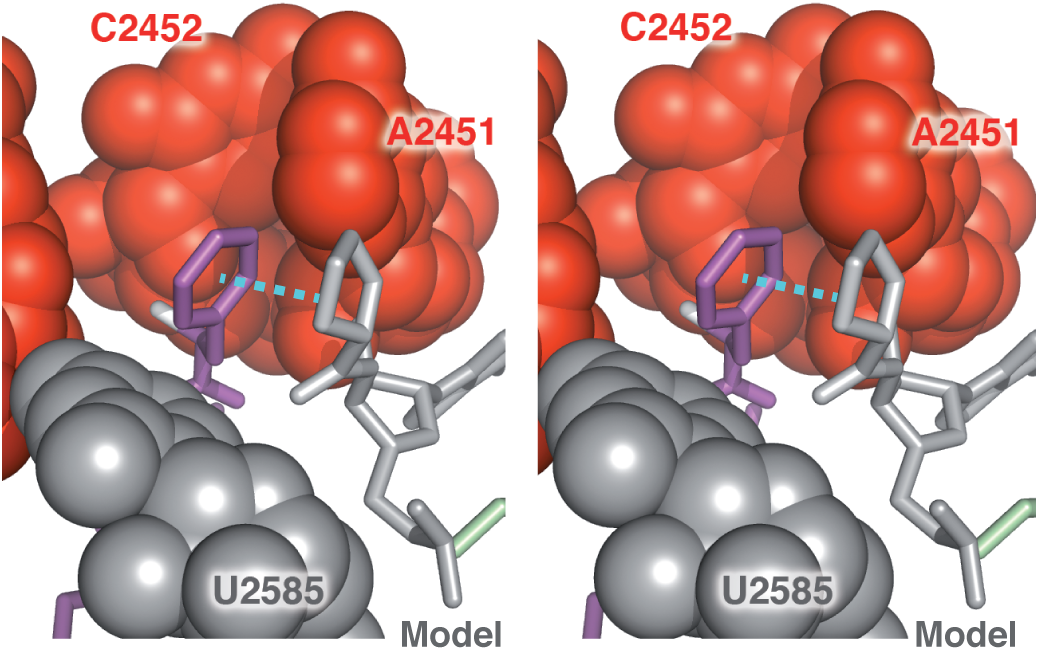
Energy minimized Psi(pi) rotamer of A-Phe 5’p aligned in the PTC cavity (from pdb 1VQ6, as in Figure 5A), with proline as P-site residue (stereo). Residues in grey are from pdb 2WDM (P-site A-Phe mutated to A-Pro) and pdb 2WDN (orientation of U2585). The ring-ring distance in this (non-optimized) Phe-Pro configuration (light blue dotted line) is ~3.9 Å, close to the maxima of the distribution established in proteins (Brandl et al. 2001; Bhattacharyya and Chakrabarti 2003). A-site aminoacyl tRNA is in purple; P-site aminoacyl tRNA is in green.

**Figure S2.**
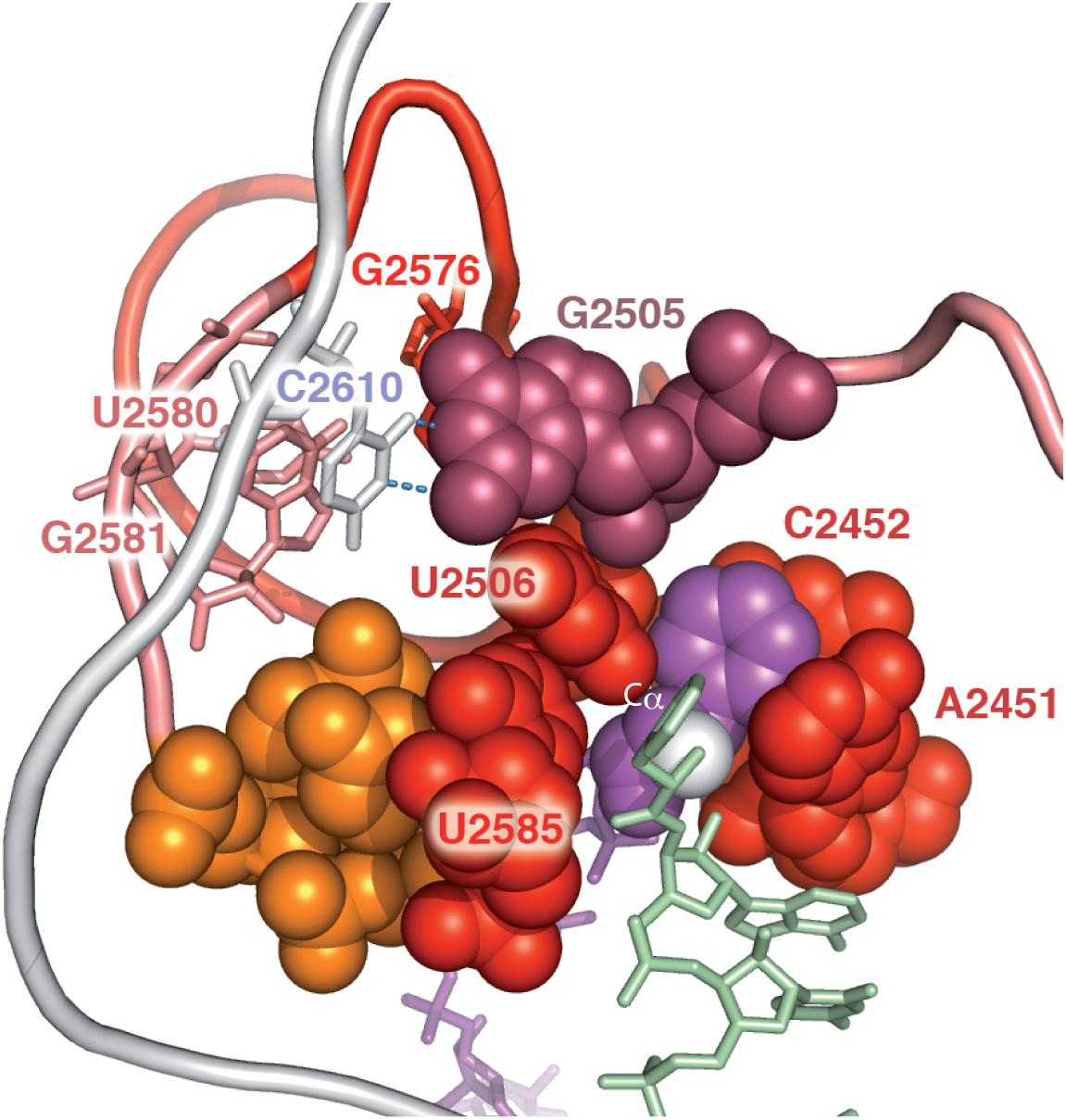
On the rationale of the PTC architecture: overview of the induced PTC, highlighting ribosomal residues involved in the mechanism orienting the A-site aminoacyl ester by induction. In order for U2506 to enforce the reactive Psi rotamer through a direct contact with the Cα atom, movements of this residue from the induced position must be prevented. The PTC is organized in such a way as to firmly keep the G2505 nucleotide in a stretched orientation that, in conjunction with U2585, holds U2506 in place. The stability of G2505 is achieved through a reversed WC base pair with C2610, which in turn stacks on G2581 and U2580. In addition, G2505 optimally stacks on G2576 (behind G2505 in the Figure). The requirement for a guanine at this location generates a peculiar arrangement of the strand that places G2576. A-site Phe(N) is shown in white. Unmodified pdb 2WDM and 2WDN structures (Voorhees et al. 2009).

**Figure S3.**
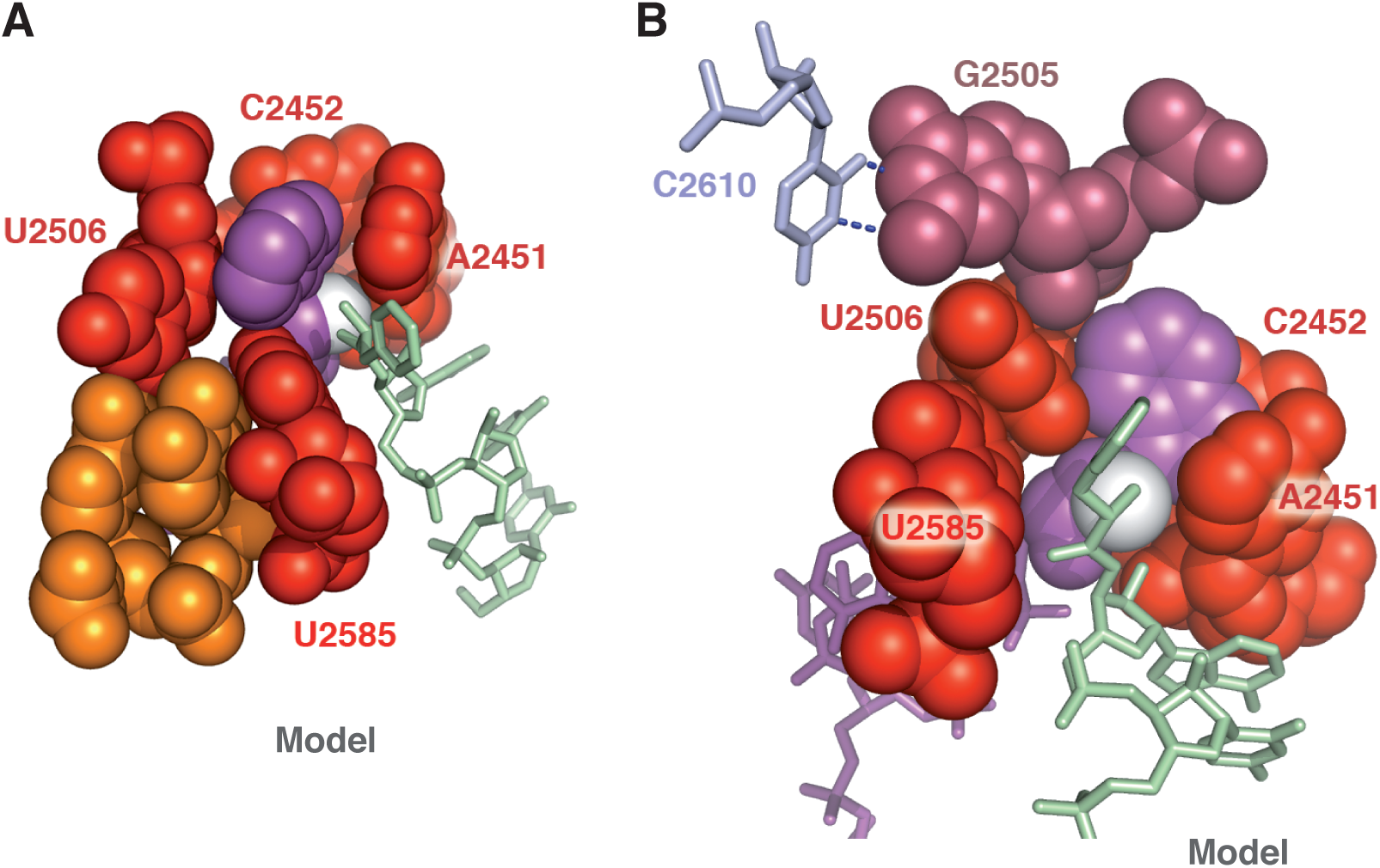
PTC before (A) and after (B) induction, with tryptophane as the A-site aminoacyl ester. (A) Uninduced state. The confinement ofthe si de-chain revealsthattee entering of tryptophanyl ester inside the PTC cavity requires a substantial displacement of U2585, shown here in its actual orientation in pdb 1VQ6 (Schmeing et al. 2005). (B) After induction, the side-chain is trapped inside the peptidyl transferase center cavity (from pdb 2WDM and 2WDN, Voorhees et al. 2009). A-site aminoacyl tRNA is in purple; P-site aminoacyl tRNA is in green. In A and B, A-site Trp(N) (the nucleophile) is shown in white. These configurations were obtained with an A-site Phe → Trp mutation in the pdb files as the only modification (see method Section).

**Text S1: Additional considerations on the volume correlation in the genetic code and its kinetic interpretation**

The correlation is shown in (Lehmann 2000, Fig. 3). It was interpreted in the structural and kinetic context shown in (Lehmann 2000, Fig. 5; Lehmann et al. 2009, Figs. 1 and 4). Our interpretation of this correlation is based on some hypotheses that we wish to further clarify here.

1. A direct connection between anticodon–codon free-energy-change interactions (ΔG_0_ ac-c) and the dissociation rate constant ***k_-_*** is made under the hypothesis that the association rate constant ***k***_+_ is uniform among different complementary anticodon-codon interactions (Lehmann 2000; Lehmann et al. 2009). With ΔΔG_0_ ac-c max ≈ 4 kcal mol^−1^ (see the correlation), the ratio between the highest and the lowest ***k****_-_* is ~1000.
2. To the best of our knowledge, there is no available data that may allow an assessment of the influence of the side-chains of the aminoacyl esters on the kinetics of peptide bond formation in a context without local confinement. However, data from intramolecular systems (Bruice and Pandit 1960; Lightstone and Bruice 1996) suggest that for a given backbone configuration (Lightstone and Bruice 1996: compounds I, II, III, V and VI in Table 1), the ratio between the highest (compound VI) and the lowest (compound I) rate constant ***k_rel_*** is ~1000. A similar ratio between the extreme ***k_cat_*** values of the amino acids could thus also characterize the (unknown) reactional context on the early ribosome. A major difference between these intramolecular systems and the aminoacyl esters is however the level of *gem* substitution (disubstitution vs. monosubstitution).
3. Combining (1) and (2), the possible correspondence between Δ***k_-_*** max and **Δ*k_cat_*** max is consistent with the correlation, which corresponds to a situation for which the equality ***k_cat_*** ≈ ***k_-_*** is verified for all [amino acids I codon] couples (Lehmann 2000; Lehmann et al. 2009). Conversely, *the correlation* suggests that **Δ*k_cat_*** max is ~1000, in possible agreement with the data of (Bruice and Pandit 1960).
4. The structural context of the reaction of peptide bond formation associated with the mentioned correlation is seemingly different from the context of the uninduced state of the PTC. The correlation suggests a *monotonic* increase of *k_cat_* with the size of the side-chains, a behavior that is clearly not observed in Fig. 1 (e.g. considering the sequence A-Gly, A-Ala, A-Val, A-Phe). This monotonic property is typically (albeit not always) observed in intramolecular systems (see Lightstone and Bruice 1996; Jung and Piizzi 2005), for which the kinetics is established in solution (i.e. without confinement). This similarity suggests to us an absence of catalytic site on the early ribosome, at least a catalytic site where reactive species are confined. As exemplified by the A-Gly acceptor, residues in the PTC can dramatically perturb the orientation of the nucleophile in the uninduced state (Fig. 1 & Fig. 4), resulting in an almost complete loss of activity with this amino acid (Rychlík et al. 1970), a situation that the correlation does not reflect. It thus appears that the PTC architecture responsible for the confinement mechanism could not occur without some “inconvenience” for several aminoacyl esters in the uninduced state, in which U2585 can dramatically reduce the activity of some A-aa. From the point of view of bacteria, this “inconvenience” could be seen as an “advantage” since only few types of A-aa (and the Pm analogue) may play the role of reactive antibiotics. There are three outliers in the volume correlation: Asn, Arg and Trp. The situation for Arg and Trp suggests a discontinuity related with size, which recalls an effect discussed in the context of the PTC (see discussion Section). Innovative experiments could unravel the reason for this similarity.
5. A major question is whether the early ribosome had any “pre-defined” catalytic site at all. An absence of catalytic site imply that the role of the early ribosome boiled down to maintaining peptidyl-tRNA and aminoacyl-tRNA juxtaposed during translation, with no confinement of the aminoacyl moiety. According to our analysis, this implies a “side-chain effect”, the optimization of which results in the volume correlation (Cibils *et al*., work in preparation).

## Footnotes

2 Lightstone and Bruice (1996): Table 1; Jung and Piizzi (2005): Tables 1 and 4.

